# dsRBD Redesign: A Targeted Strategy for Inhibition of RNA Helicase DHX9

**DOI:** 10.64898/2026.02.13.705762

**Authors:** Nina Lang, Emily Freund, Lucie Haene, Kristian Schweimer, Janosch Hennig

**Affiliations:** Department of Biochemistry IV – Biophysical Chemistry, University of Bayreuth, 95447 Bayreuth, Germany; Molecular Systems Biology Unit, European Molecular Biology Laboratory (EMBL), 69117 Heidelberg, Germany

## Abstract

The RNA helicase DHX9 is essential for genomic stability, transcription, translation regulation and other RNA-related processes. DHX9 has emerged as a therapeutic target for cancer treatment as its expression levels are elevated in several different cancer types. Moreover, tumor cells exhibit a strong dependence on DHX9, making its inhibition an effective strategy for tumor regression. As RNA helicases are conserved enzymes, unique features need to be targeted to minimize side effects. Here, we identified an autoregulatory interface between the DHX9 helicase core and double-stranded RNA binding domain 2 (dsRBD2) ideal for a highly specific inhibition of DHX9. By targeting the proposed dsRBD2-core interface in DHX9, we aim to specifically inhibit DHX9 helicase activity by preventing dsRBD2 binding to the helicase core. We developed a protein binder design-based strategy targeting the dsRBD2-core interface of DHX9 by computationally designing novel dsRBDs. By binding this interface without engaging RNA, the dsRBD designs prevent dsRBD2-core interaction and inhibit DHX9 helicase activity. Our strategy of redesigning autoregulatory protein domains as inhibitors offers a computationally efficient alternative to larger-scale library generation and provides a flexible framework applicable to a wide range of therapeutic targets.

## Introduction

Cancer development is a complex process involving genetic mutations, epigenetic alterations, and dysregulation of key cellular pathways by oncogene activation or tumor suppressor gene inactivation^1,2^. Tumorigenesis is often driven by dysregulated gene expression, abnormal RNA processing, and increased protein synthesis, resulting in malfunctioning of key regulatory pathways^3–5^. Consequently, cancer cells rapidly grow and evade regulatory mechanisms. In this context, RNA helicases have been shown to be crucial regulators of tumorigenesis as they remodel RNA structures and are involved in nearly all aspects of RNA metabolism^6,7^. In human cells, they contribute to all RNA-related cellular processes such as transcription, translation, pre-mRNA splicing, rRNA processing or RNA degradation^8–10^. Thus, precise regulation and activity of RNA helicases is crucial for maintaining cellular homeostasis. Aberrant RNA helicase activities including overexpression, hyperactivity or functional defects have been associated with cellular stress and linked to various diseases, including cancer development, neurological disorders, and developmental abnormalities^10–13^. Given their central role in keeping RNA metabolism in balance, RNA helicases have emerged as potential biomarkers and therapeutic targets in cancer research.

RNA helicases are a diverse class of enzymes that utilize the energy of nucleoside triphosphate (NTP) hydrolysis to unwind diverse RNA structures and remodel RNA-protein complexes^14–16^. The widespread distribution of RNA helicases across bacteria, archaea, eukaryotes and viruses highlights their fundamental importance.

The RNA helicase DHX9, also known as RNA helicase A (RHA) or nuclear DNA helicase II (NDHII) is a DEAH/RHA helicase and has been shown to be associated with tumorigenesis. DHX9 is a functionally diverse helicase as it acts on RNA and DNA as well as on hybrid substrates or triple-helical DNA structures^17–20^. Furthermore, DHX9 is ubiquitously expressed and plays a critical role in maintaining genomic stability by regulating transcription, translation, RNA processing including splicing and editing, RNA transport, miRNA genesis, DNA repair and is involved in the formation of intronic RNA containing stress granules^21–25^. Due to its regulatory roles, DHX9 has been shown to be a key regulator in several different cancer types^22,26–32^. More specifically, overexpression and thus elevated DHX9 activity is observed in lung, prostate, cervical or colorectal cancer, as well as in Ewing sarcoma, myelodysplastic syndromes and multiple other cancer types^22,23,31,33–39^. Often DHX9 upregulation is associated with increased malignancy and poor prognosis of patients^40^. Furthermore, elevated DHX9 activity enhances DNA damage repair, suppresses apoptosis, and modulates gene expression, making cancer cells more resistant to chemotherapeutic drugs such as cisplatin or carboplatin^41^. Additionally, elevated DHX9 activity is associated with key hallmarks of cancer, including gene regulation, sustained proliferative signaling, evasion of growth suppressors, resistance to apoptosis, angiogenesis, and metastasis^42^.

Given its multifaceted roles, DHX9 is a potential therapeutic target in cancer treatment. Hence, there is the need to develop specific DHX9 inhibitors as a potential therapeutic strategy in DHX9 responsive diseases and cancer types. Despite DHX9 activity is essential during early development in mammals, a systemic knockout in adult mice is well tolerated, which enables effective targeting in human cells^43^. Furthermore, different types of cancer cells depend on DHX9 activity as depletion results in tumor regression^43^. Given the suitability of DHX9 as a therapeutic target, strategies need to be developed to specifically target DHX9. However, since DHX9 is one of many RNA helicases with conserved domains, making selective targeting essential to avoid off-target effects. The conserved helicase core includes two RecA-domains, which mediate nucleoside triphosphate (NTP) binding and hydrolysis. Thus, targeting the ATP-binding pocket or other conserved domains is not suitable for specifically inhibiting DHX9 as this would disrupt the function of multiple RNA helicases within the cell. Therefore, drug strategies should focus on unique features of DHX9 to enhance specificity and efficiency with minimal off-target effects as this will achieve greater precision towards DHX9, reduce toxicity and improve treatment outcomes. A recent study has identified a small molecule that inhibits DHX9 which binds in a pocket located in RecA1 nearby the exit of the RNA tunnel, thereby inhibiting RNA unwinding^44^. Despite its conserved helicase core, DHX9 contains unique auxiliary domains consisting of two N-terminal double-stranded RNA binding domains (dsRBDs) and a disordered C-terminal RGG-rich region, offering potential selective target sites. While structural data is available for DHX9 bound to ADP showing that dsRBD2 is flexible in this state, more is known about its *Drosophila melanogaster* ortholog maleless (MLE)^45,46^. Cryo-EM studies of MLE capturing different conformational states revealed that MLE-dsRBD2 regulates helicase activity by aligning dsRNA and interacting with the helicase core, an interface critical for function^46^. Given the 51% identity and 85% similarity between MLE and DHX9, we hypothesize that the DHX9 core-dsRBD2 interaction may serve a similar regulatory role.

Here, we present a protein-based strategy targeting the dsRBD2-core interface for RNA helicase inhibition. Combining different approaches allows for targeting multiple functional sites, thereby improving treatment efficacy and can help to overcome resistance. Thus, we target the highly specific dsRBD2-helicase core interface by redesigning dsRBD2 using a computational approach. First, we generated a pool of new dsRBD sequences with dsRBD2 as input and sequence restraints using ProteinMPNN^47^. Using AlphaFold2 and AlphaFold3 the structures of the generated sequences were predicted and filtered for the dsRBD-fold^48–50^. Furthermore, our designs were docked to the helicase core using HADDOCK^51,52^. The four most promising designs were then experimentally tested. Three of the four tested designs adopt the expected dsRBD-fold, whereas one displays mispacking of secondary structure elements, suggesting an incomplete or unstable fold. Interaction studies between the isolated helicase core domain (DHX9core, aa 268-1148) and dsRBD2 and helicase inhibition assays further support the potential of our designs to function as DHX9 inhibitors. Taken together, we present a novel interface for targeted DHX9 inhibition with potential applications in future drug development. In addition, our strategy to target autoregulatory domains using redesigned dsRBDs introduces a new approach to modulate DHX9 activity and could likely be expanded to other systems with autoregulatory functions. Furthermore, our strategy displays a proof-of-principle from which potent peptide-like inhibitors can be derived for targeted DHX9 inhibition.

## Results

### Functional importance of the dsRBD2-core interface for DHX9 activity

DHX9 and its ortholog in *Drosophila melanogaster*, MLE, are both nuclear RNA helicases involved in RNA processing and gene regulation. While MLE is well-studied in *Drosophila* in the context of dosage compensation and remodeling of roX2 lncRNA, DHX9 is involved in several different RNA-related processes. Their high sequence similarity suggests a similar enzymatic mechanism (Figure 1a). Recently, a novel interface in MLE between dsRBD2 and the helicase core was shown to be functionally relevant for helicase activity. In fact, dsRBD2 acts as an autoregulatory unit and is not only recruiting dsRNA to the RNA tunnel but also required to open the RNA tunnel by specifically binding to the helicase core while contacting RecA2, OB-fold and HA2 domain (Figure 1b,c)^45,46^. This interface is thus crucial for helicase activity and presents itself as a novel target site for drug development. To date, structural information on full-length DHX9 remains limited, as available structures only cover the conserved helicase core excluding the auxiliary domains dsRBD1, dsRBD2 and the C-terminal RGG-rich region. However, AlphaFold2 predicts an interaction between dsRBD2 and the helicase core resembling that of MLE, whereas dsRBD1 is modeled without specific interactions to other parts of DHX9 (Figure 1d). Therefore, we hypothesize that dsRBD2 might exhibit a similar regulatory mechanism in DHX9 as shown for MLE with the dsRBD2-core interface being crucial for helicase activity. To assess whether a protein interacting at this interface modulates helicase activity, several complementary approaches can be used. First, the DHX9_core_ interface can be mutated to disrupt binding of the protein. Changes in helicase activities would indicate the functional relevance of the interaction at the interface. Second, a competing protein can be added while monitoring helicase activity. Competition at the binding interface will result in a change in helicase activity indicative of a functional relevant interface. Here, we supplemented wildtype dsRBD2 as an inhibitor and monitor helicase activity of DHX9.

**Figure 1:**
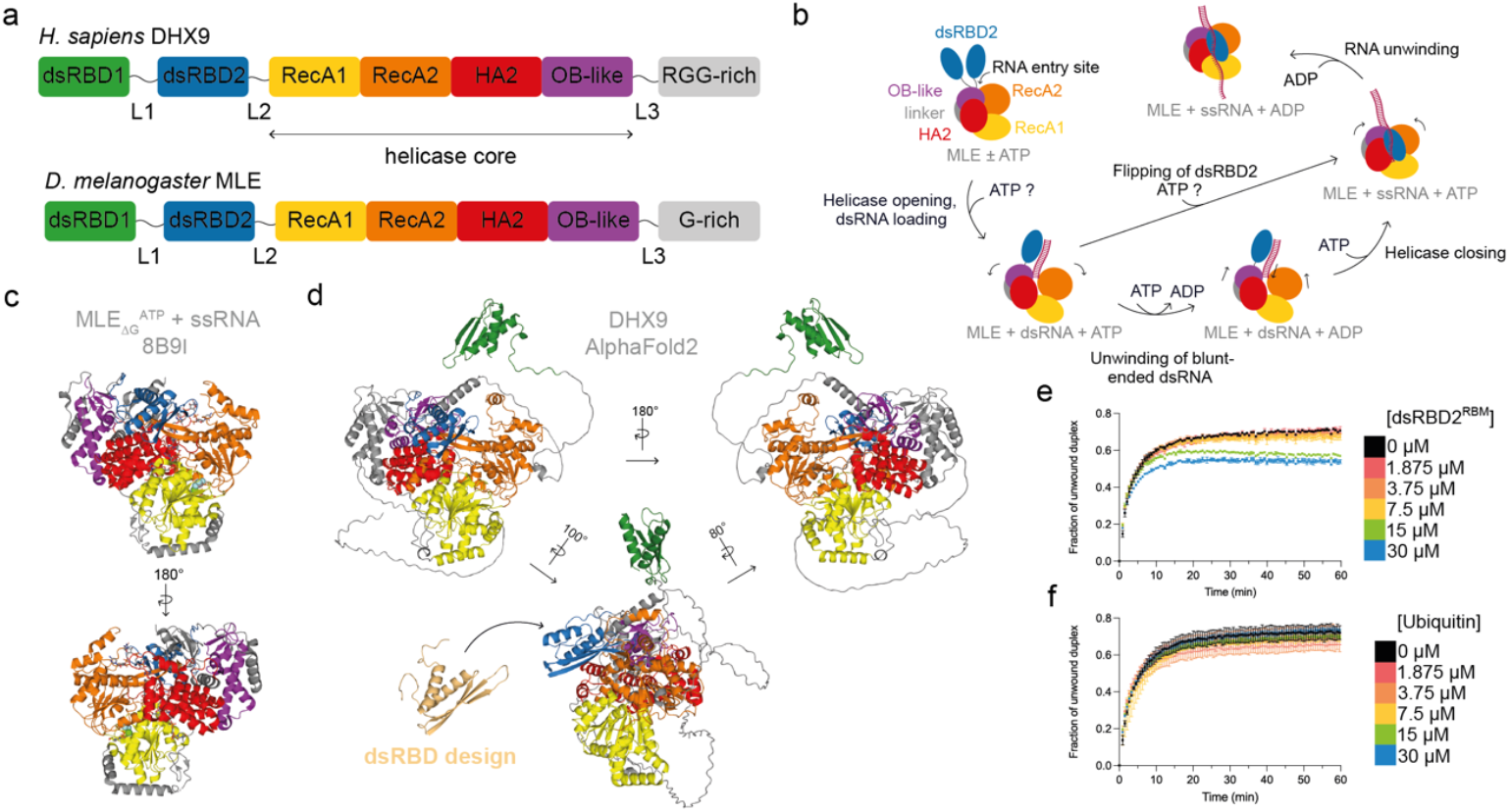
The helicase mechanism of MLE and its human ortholog DHX9 proposes a novel target site for DHX9 inhibition. **(a)** Conserved domain organization of *H. sapiens* DHX9 and *D. melanogaster* MLE. The region spanning the helicase core is indicated for DHX9. **(b)** MLE RNA helicase mechanism^9,46^. Domain coloring is according to panel a. **(c)** Cryo-EM structure of MLE _ΔG_ in complex with ssRNA showing the functional important dsRBD2-core interface. PDB: 8B9I^46^. Domain coloring is according to panel a. **(d)** AlphaFold2 model of DHX9 showing the predicted dsRBD2-core interface, while dsRBD1 is predicted to not interact with other parts of DHX9 is likely freely tumbling in solution in absence of other binding partners. The target site for our designs is competing with the dsRBD2 binding site and is indicated with the dsRBD design. **(e)** Helicase assay of DHX9 with different concentrations of a dsRBD2 RNA-binding mutant (dsRBD2^RBM^) showing the potential to function as a DHX9 inhibitor. **(f)** Helicase assay of DHX9 with the addition of ubiquitin as a negative control showing no effect on helicase activity.

To determine whether the supplementation with dsRBD2 can inhibit DHX9 helicase activity, we generated an RNA-binding-deficient dsRBD2 mutant (dsRBD2^RBM^) using previously reported mutations thereby isolating the effects of protein-protein interactions with the helicase core from those of RNA binding^53^. Notably, helicase activity of DHX9 is slightly downregulated in the presence of dsRBD2^RBM^ in a concentration dependent manner (Figure 1e). To exclude a concentration dependent crowding effect of the additional protein in the reaction, we performed the same helicase assay with ubiquitin, which should not interact with DHX9. Accordingly, addition of ubiquitin has no effect on helicase activity (Figure 1f). Thus, dsRBD2^RBM^ is binding to the DHX9_core_ interface competing with native dsRBD2, thereby preventing native dsRBD2 from binding and reducing helicase activity (Figure 1d). This specific region within the helicase core reveals a novel therapeutic target site for DHX9. Furthermore, the dsRBD-fold displays a promising protein scaffold for targeted binding and inhibition.

### Design of novel dsRBDs derived from DHX9 dsRBD2

To design potential inhibitors, we chose a protein binder design approach based on redesigning a dsRBD aimed to target the proposed dsRBD2-core interface to specifically inhibit DHX9. Instead of simply using wildtype dsRBD2 or a shortened peptide we designed new sequences derived from dsRBD2 using ProteinMPNN^47^. Through redesigning dsRBD2 we can control the properties and interactions of the molecule, ensuring higher stability and specificity. To create a potential design for a proteinaceous DHX9 inhibitor, we developed a combined workflow that uses both machine learning tools and rational design based on dsRBD2 (Figure 2a).

**Figure 2:**
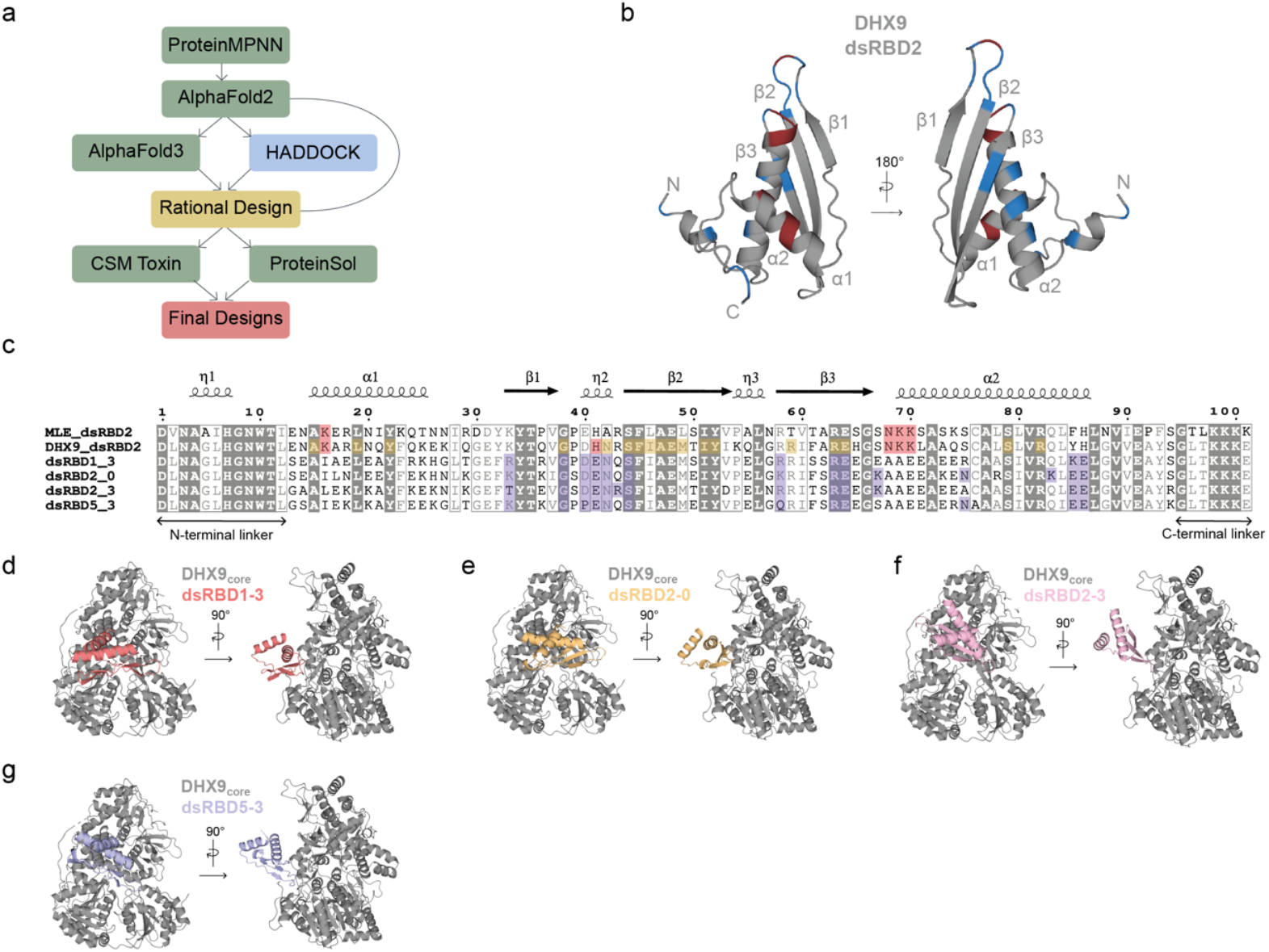
Generation of novel dsRBD designs. **(a)** Computational design pipeline novel dsRBDs based on dsRBD2. **(b)** AlphaFold2 prediction of DHX9 dsRBD2. RNA binding residues are shown in red, while residues predicted to be involved in the helicase core interaction, based on AlphaFold2 and AlphaFold3 models, are displayed in blue. **(c)** Sequence alignment of MLE dsRBD2, DHX9 dsRBD2 and the four experimentally characterized dsRBD designs dsRBD1-3, dsRBD2-0, dsRBD2-3 and dsRBD5-3. RNA binding residues are shown in red and were mutated in our designs into alanines, glutamic acids or (iso)leucines. Fixed residues are shown in yellow, which are a combination of potential important helicase core-interacting residues and conserved residues. Residues predicted to be involved in design-core contacts based on AlphaFold3 complex predictions are shown in purple. **(d-g)** HADDOCK models of dsRBD1-3 (d), dsRBD2-0 (e), dsRBD2-3 (f) and dsRBD5-3 (g) bound to DHX9_core_ (see Table 1 for docking statistics).

During our design process we mutated known RNA-binding residues located in α1 helix, β1-β2 loop and α2 helix including the conserved lysines to non-polar alanines, negatively charged glutamic acids or hydrophobic (iso)leucines to prevent RNA binding (Figure 2b,c)^53^. We analyzed sequence alignments of DHX9-dsRBD2, MLE-dsRBD2 and other dsRBDs to identify conserved residues and identified contacts between MLE-dsRBD2 and the MLE-helicase core bound to ATP-analog and ssRNA (Figure S1a). These, we compared with predicted residues involved in protein-protein interactions between DHX9-dsRBD2 and the helicase core – more specifically RecA2, OB-fold and HA2 domain – based on AlphaFold2 predictions of DHX9 with and without ssRNA (Figure S1b,c). All potential dsRBD2 residues involved in the DHX9_core_ interaction are highlighted in Figure 2b. Based on structural data known from MLE and DHX9 models it is likely that dsRBD2 mainly interacts with the helicase core with the three β-strands, while the RNA binding residues located in α1 helix, β1-β2 loop and α2 helix are still available for RNA binding. For the generation of our dsRBD designs we fixed selected residues based on a combination of potentially important residues involved in interactions with the helicase core and residues conserved among dsRBDs (Figure 2c). These residues are located within α1 helix (A15, L19, Y22), in the β1-β2 loop (G38), in β2 (S44, F45, I46, A47, E48, M49), in β3 (R59, R63, E64) and in α2 helix (S79, R82). The sequence of DHX9-dsRBD2 was then redesigned using ProteinMPNN and subsequently predicted with AlphaFold2^47–49^.

**Table 1.** Computational statistics of the dsRBD designs. ^a^AlphaFold3 interface predicted template modeling score.^b^HADDOCK root mean square deviation of the distance between the interface backbone atoms and the lowest energy structure of the same cluster. ^c^HADDOCK buried surface area. ^d^according to lowest HADDOCK score.

|  | dsRBD1-3 | dsRBD2-0 | dsRBD2-3 | dsRBD5-3 |
| --- | --- | --- | --- | --- |
| SeqNovelty | 47 % | 51 % | 45 % | 49 % |
| Fixed residues | 24 % | 24 % | 24 % | 24 % |
| *De novo* residues | 61 % | 67 % | 60 % | 65 % |
| Conserved residues | 29 % | 25 % | 31 % | 27 % |
| AF3 iptm<sup>a</sup> | 0.51 | 0.73 | 0.64 | 0.45 |
| Haddock score [AU] | -108 | -93 | -103 | -75 |
| Haddock RMSD [Å]<sup>b</sup> | 0.47 | 0.74 | 0.56 | 0.48 |
| Haddock BSA<sup>c</sup> [Å<sup>3</sup>] | 2317.9 | 2008.4 | 1862.8 | 1492.6 |
| No. of clusters | 12 | 15 | 12 | 12 |
| No. structures in best cluster<sup>d</sup> | 34 | 20 | 22 | 48 |
| CSM Toxin | Non-toxic | Non-toxic | Non-toxic | Non-toxic |
| Protein Sol | 84.4 % | 84.8 % | 97.3 % | 93.2 % |

Out of 30 designed sequences, 12 sequences (40%) passed our pLDDT standards (≥ 80). The 12 models were further analyzed in their compatibility in binding to DHX9_core_ at our desired dsRBD2-core interface using AlphaFold3 and HADDOCK^50–52^ (Figure S1a). We tested them for toxicity and solubility probabilities using CSM toxin and ProteinSol, respectively (Table 1 and Table S1)^54,55^. The designs dsRBD4-4 and dsRBD3-2 did pass our initial pLDDT standards (≥ 80) but are not predicted to bind at the dsRBD2-core interface are were therefore not carried further (Figure S1b,c). Four final designs, namely dsRBD1-3, dsRBD2-0, dsRBD2-3 and dsRBD5-3 are predicted to bind to the core and were further selected for biochemical and biophysical characterization (Figure 2d-g and S1d-g). The sequence novelty of these four designs was assessed using SeqNovelty^56^. Despite having 24 % fixed residues, our final designs had over 60 % *de novo* residues, resulting in SeqNovelty scores ranging from 45 % to 51 %. Among these, dsRBD2-0 and dsRBD5-3 exhibit the highest novelty (Table 1).

### dsRBD designs are folded, thermostable and RNA-binding deficient

All four designs expressed soluble and can be purified with high purity as validated by the polydispersity index (PDI) using dynamic light scattering (DLS, Figure 3a). Analysis of the hydrodynamic diameter shows similar diameters compared to DHX9 dsRBD2, while dsRBD1-3 shows a slightly increased diameter, indicating looser packing or slight aggregation of the design (Figure 3a,b). Remarkably, all our designs exhibit high thermal stability with melting temperatures ranging from 76.9 °C up to 89.0 °C exceeding the melting temperature of DHX9-dsRBD2 by 14-26 °C (DHX9-dsRBD2: 62.6 °C, Figure 3c). To evaluate the folding properties of our designs, we recorded CD spectra that reveal a secondary structure content similar to DHX9-dsRBD2 with a mixture of α-helices and β-strands, as indicated by a maximum around 195 nm, minima at 208 nm and 222 nm reflecting contributions from both secondary structure elements (Figure 3d). The folding is further confirmed by 1D NMR spectra showing disperse amide regions between 6 ppm and 8.5 ppm. However, dsRBD1-3 again seems to be less structured as the regions below 6 ppm and above 10 ppm are less dispersed and have lower signal to noise ratio (Figure 3e. Methyl regions of the designs are well dispersed and show peaks upfield of 0.5 ppm indicative of folded proteins and higher-order structure (Figure 3f). Importantly, our designs do not interact with either dsRNA or ssRNA, eliminating RNA-binding interference and enabling reliable analysis in helicase activity assays involving RNA (Figure 3g,h).

**Figure 3:**
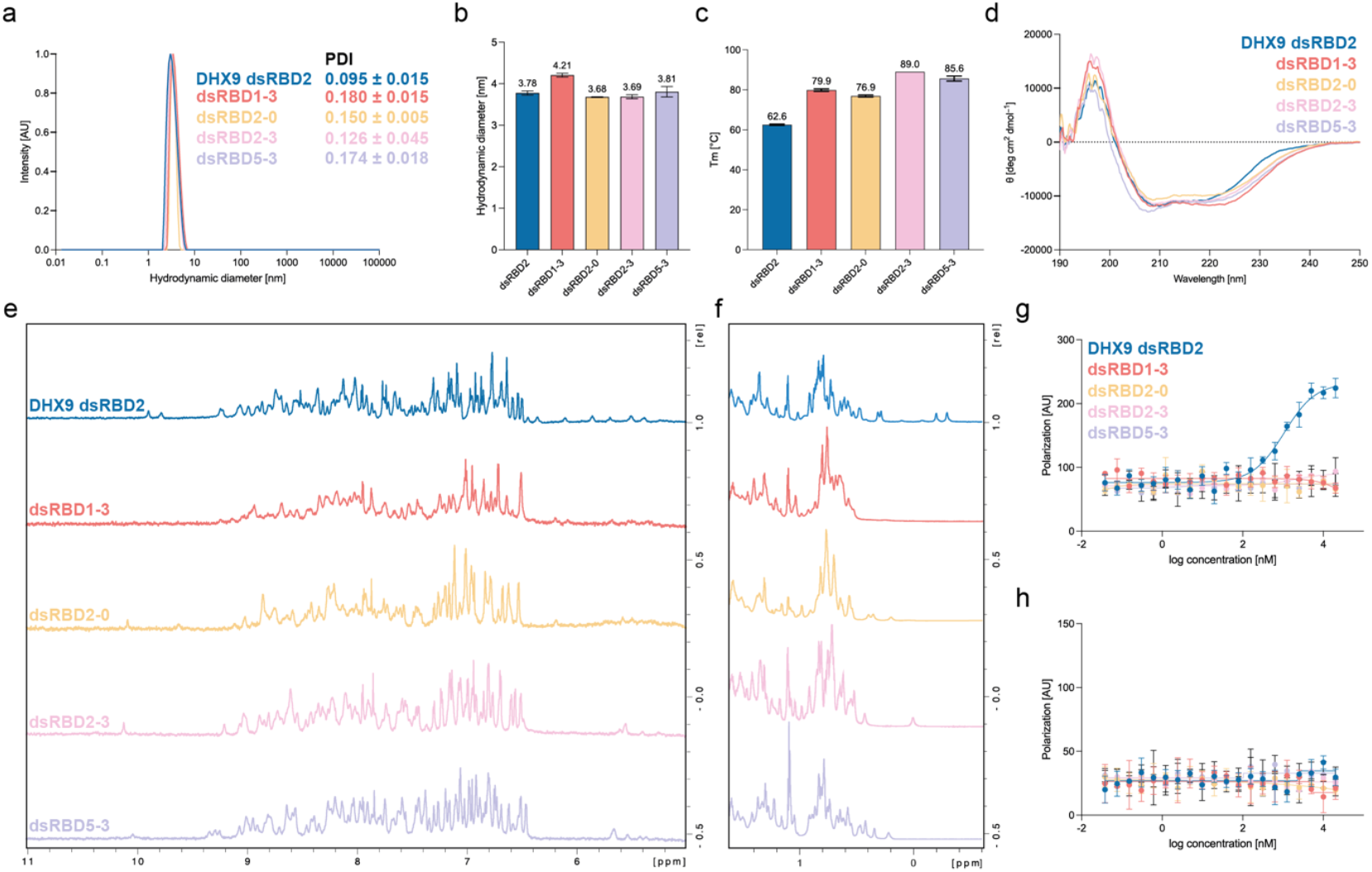
Biochemical and biophysical characterization of the dsRBD designs. **(a)** DLS data of the dsRBD designs showing the normalized intensity of the hydrodynamic diameter and the polydispersity index (PDI). For each protein one representative DLS curve is shown. **(b)** DLS data showing the hydrodynamic diameter. Measured in triplicates. **(c)** Melting temperatures of the dsRBD designs determined with the barycentric mean. Measured in triplicates. **(d)** Circular dichroism spectra of the dsRBD designs. **(e)** 1D NMR spectra showing the proton amide region (5-11 ppm) of the dsRBD designs. (f) 1D NMR spectra showing the methyl region (−0.5-1.5 ppm). **(g and h)** Fluorescence polarization assays with dsRNA (g) and ssRNA (h). Assays were conducted in triplicates. DHX9 dsRBD2 is used as a positive control.

### In-depth structural analysis reveals differences between designs

So far, all designs exhibited similar biochemical properties, although dsRBD1-3 appeared to contain secondary structure elements that might be poorly packed in the 3D structure, possibly due to weak intramolecular interactions between the secondary structure elements. This is further supported by the analysis of ^1^H, ^15^N HSQC spectra of the designs as the signals of the amide proton resonances show less dispersity in dsRBD1-3 being mainly between 8 and 9 ppm indicative of partially unstructured regions (Figure 4a). Instead, ^1^H, ^15^N HSQC spectra of dsRBD2-0, dsRBD2-3 and dsRBD5-3 show dispersed amide proton resonances between 7 and 10 ppm characteristic for well-folded proteins (Figure 4b-d). Thus, we focused on dsRBD2-0, dsRBD2-3 and dsRBD5-3 and performed protein backbone assignments by standard triple resonance NMR spectroscopy. Analysis of the secondary chemical shifts reveals the presence of the typical α-β-β-β-α fold of dsRBDs and further confirms the AlphaFold2 predictions (Figure 4e-g)^57,58^. Interestingly, β1-strands in our designs appear to extend further toward helix α1 and is therefore longer than predicted by AlphaFold2, which likely improves packing against β2 and thus stabilizes the overall fold.

**Figure 4:**
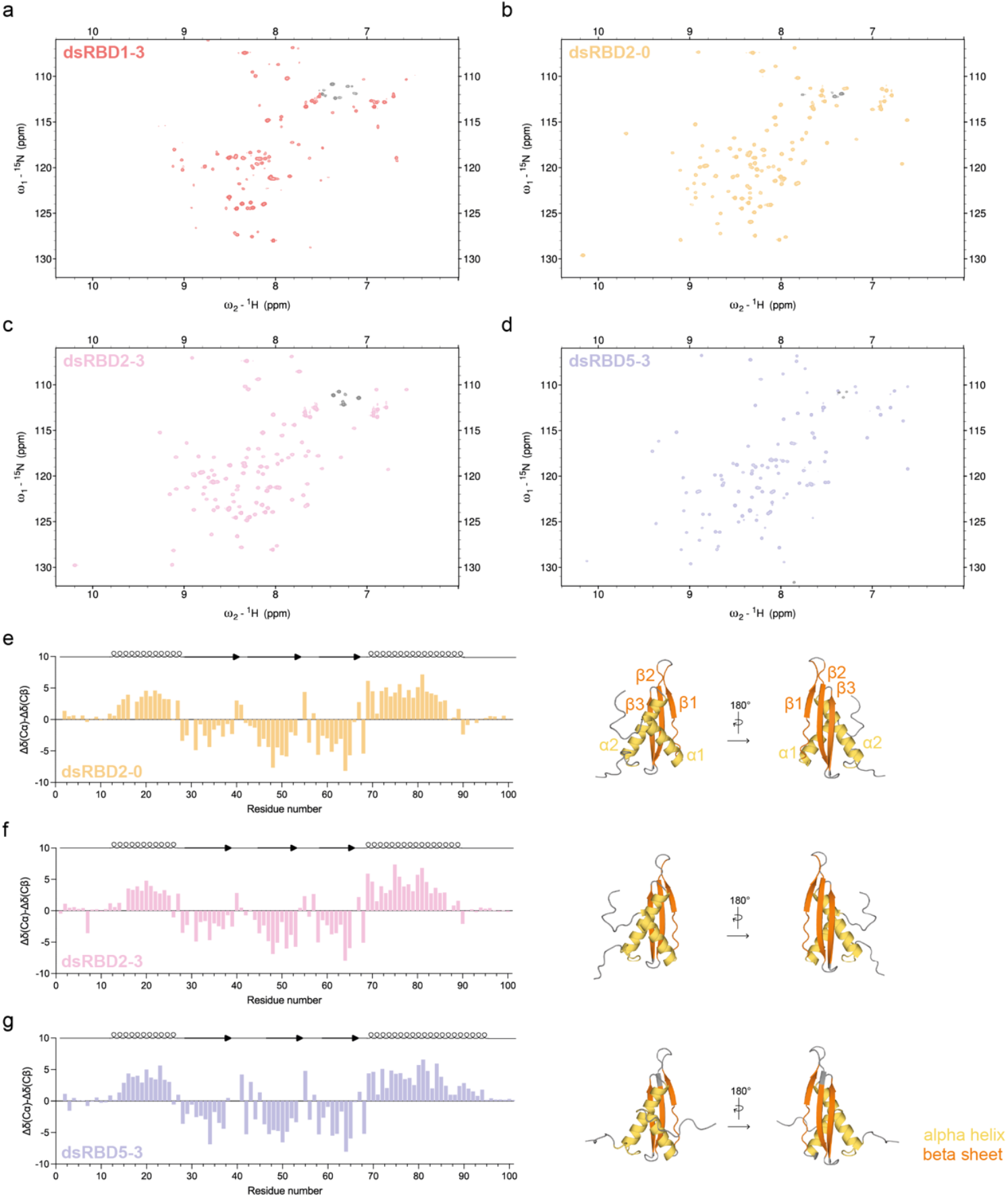
Secondary structure determination of the designs. **(a-d)** ^1^H ^15^N HSQC spectra of dsRBD1-3 (a), dsRBD2-0 (b), dsRBD2-3 (c), dsRBD5-3 (d). **(e-g)** Secondary chemical shifts of dsRBD2-0 (e), dsRBD2-3 (f) and dsRBD5-3 (g) showing the α-β-β-β-α dsRBD fold. Corresponding secondary chemical shifts are plotted on the AlphaFold2 predictions. Alpha helices are shown in yellow, beta strands are shown in orange.

To further validate the designs, we determined the solution structure of dsRBD2-0 by NMR spectroscopy (PDB accession code: 28PM). The ensemble of the 20 lowest energy structures displays low violations of experimentally derived constraints together with small deviations from ideal geometry (Figure 5a, Table 2). dsRBD2-0 shows the typical dsRBD fold with residues 12-88 forming an α-β-β-β-α fold consisting of three anti-parallel β-strands with two α-helices on one side (Figure 5b). The AlphaFold2 model aligns with the NMR structure with a Cα RMSD of 1.639 (residues 12-88) (Figure 5c). However, α1 is shifted towards the β-strands in the NMR structure compared to the AlphaFold2 model.

**Table 2:**
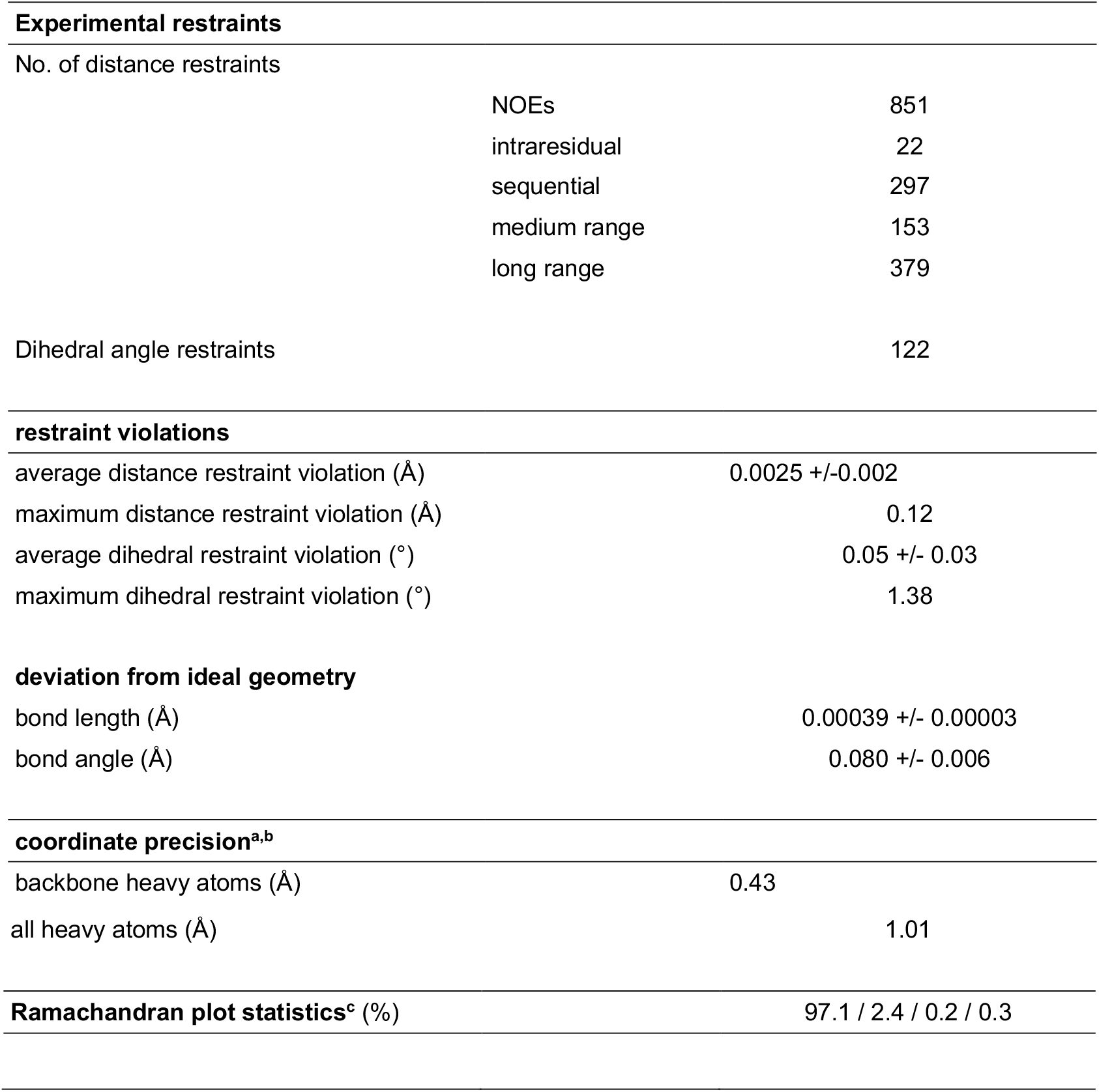
Structural statistics for the NMR solution structure of dsRBD2-0. ^a^The precision of the coordinates is defined as the average atomic root mean square difference between the accepted simulated annealing structures and the corresponding mean structure calculated for the given sequence regions. ^b^Calculated for residues 13-39, 44-92 (structured regions). ^c^Ramachandran plot statistics are determined by PROCHECK^82^ and noted by most favored/additionally allowed/generously allowed/disallowed.

**Figure 5:**
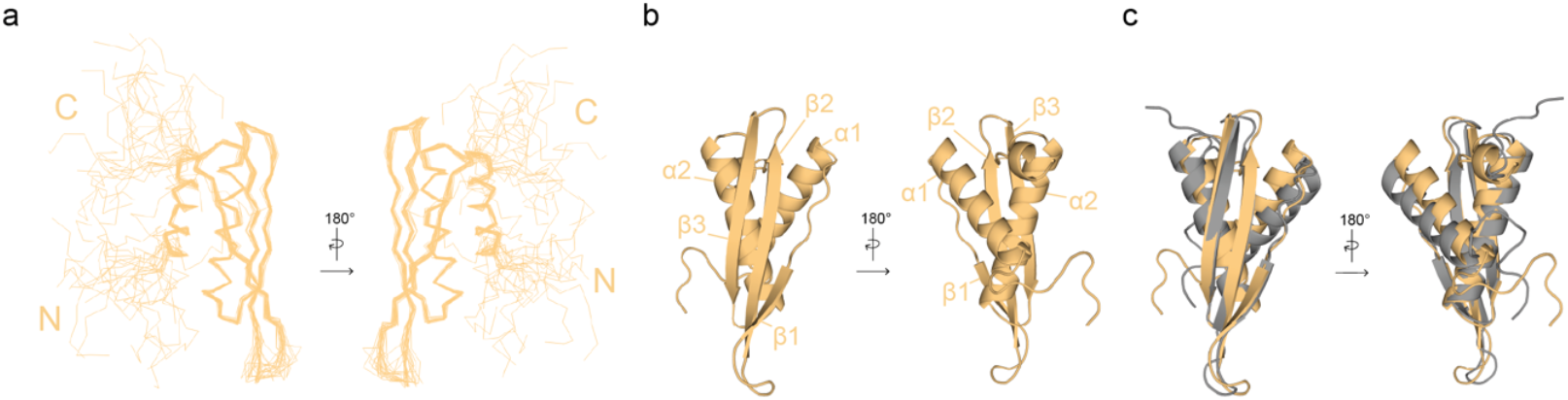
NMR solution structure of dsRBD2-0. **(a)** NMR ensemble of the 20 lowest energy structures generated by backbone superimposition of dsRBD2-0 (PDB accession code: 28PM). N- and C-termini are labelled. **(b)** Cartoon representation of the lowest energy structure of dsRBD2-0. Secondary structure elements are marked and confirm the dsRBD-fold. **(c)** Superposition of the dsRBD2-0 NMR structure (yellow) and the AlphaFold2 prediction (grey).

### Interactions of the dsRBD designs with DHX9_core_and their potential to function as DHX9 inhibitors

To determine whether our dsRBD designs interact as desired with DHX9_core_, we performed NMR titration experiments. While observing the resonances of the ^15^N-labeled dsRBD designs, we titrated DHX9_core_ at a ratio of 1:1 and determined changes of their signal intensities and chemical shift perturbations (CSPs). Interaction of the designs with DHX9_core_ increases the total molecular weight and decreases molecular tumbling, thus enhancing the transverse relaxation and consequently causing line broadening. This leads to a decrease in the signal intensity ratio between the free and bound state. As a control, we titrated DHX9_core_ to DHX9 dsRBD2. The interaction of dsRBD2 with DHX9_core_ is weak as indicated by the decrease of signal intensity to 70%. Notably, only small chemical shift perturbations, which are distributed across the structure, are observed for dsRBD2 (Figure 6a). In line with that, our data indicates that our dsRBD designs bind DHX9_core_ also with low affinity (Figure 6b-d). This is plausible given that the fixed residues were chosen based on known interaction sites of MLE-dsRBD2 and on AlphaFold2 predictions of DHX9. Furthermore, in the wild type the dsRBDs are connected with a flexible linker region, which increases the local concentration, leading to stronger interactions between both parts. However, in the context of DHX9 and MLE it is sensible that binding to the core cannot be too high, as the interaction must remain transient to permit the conformational changes, leading from helicase core binding back to dsRNA binding and vice versa, which is required for helicase activity. Across all three designs, dsRBD2-0 shows the largest decrease in signal intensity, dropping to 77% (Figure 6b). In contrast, signal intensities of dsRBD2-3 and dsRBD5-3 remain at 92%, indicating weaker interactions with DHX9_core_ (Figure 6c,d). This trend is further supported by the CSP analysis. dsRBD2-0 exhibits the largest residue-level chemical shift perturbations even stronger than native dsRBD2, whereas dsRBD2-3 and dsRBD5-3 show only minor perturbations (Figure 6b-d). However, CSPs can either be caused by direct interactions involving the observed residue or from an indirect effect caused by the change of the surrounding chemical environment due to conformational changes. Thus, from CSPs one cannot necessarily conclude that the corresponding amino acids are directly interacting with DHX9_core_.

**Figure 6:**
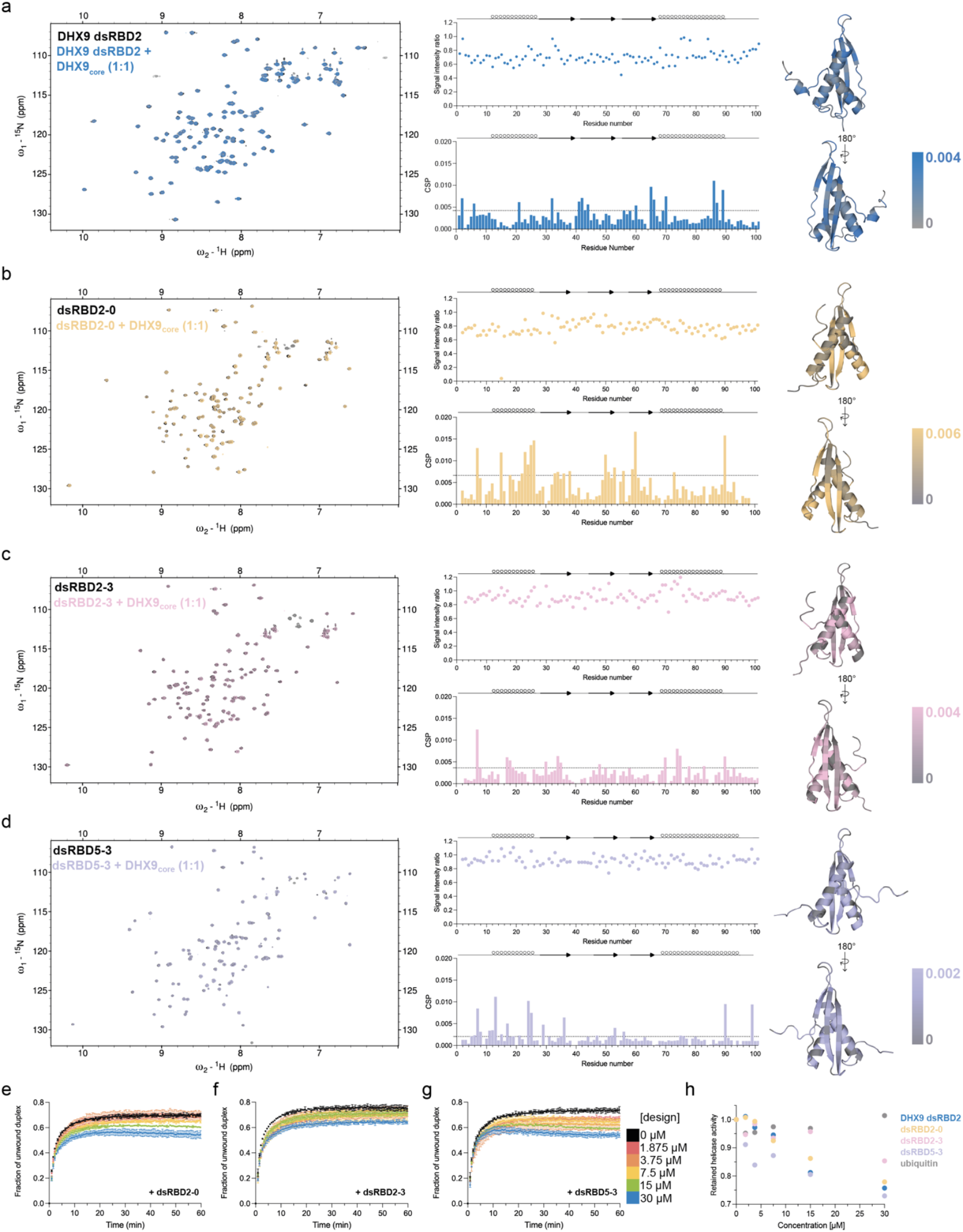
Interactions of the dsRBD designs dsRBD2-0, dsRBD2-3 and dsRBD5-3 with DHX9_core_ and their impact on helicase activity. **(a-d)** NMR titration experiments of DHX9 dsRBD2 (a), the designs dsRBD2-0 (b), dsRBD2-3 (c) and dsRBD5-3 (d) with DHX9_core_. Superimpositions of ^1^H, ^15^N HSQC spectra are shown for the free and the bound state in a 1:1 stoichiometry. Signal-intensity ratios derived from the signal intensities of the free and the bound state are plotted as well as chemical shift perturbations occurring in DHX9 dsRBD2 and the designs upon addition of DHX9_core_. Significance thresholds are indicated by the dashed line. CSPs are plotted on AlphaFold2 predictions with the maximum set to the significance threshold. **(e-g)** Helicase assays supplemented with the dsRBD designs dsRBD2-0 (e), dsRBD2-3 (f), dsRBD5-3 (g) are shown. **(h)** Plotted is the retained helicase activity depending on the design concentration in the helicase assay. The control reaction without the addition of the design was set as the reference for maximal activity.

For dsRBD2-0, the N-terminal α-helix and β_1_, β_2_ and β_3_ predominantly show CSPs, indicating conformational change upon addition of DHX9_core_ (Figure 6b). This interaction pattern suggests that dsRBD2-0 primarily contacts DHX9_core_ via the N-terminal α-helix and β-sheet, consistent with interaction modes observed for MLE-dsRBD2 and DHX9 models. These data further indicate that dsRBD2-0 binds DHX9_core_ in the intended orientation. It furthermore suggests that dsRBD2-0 might bind with greater orientational specificity compared to DHX9 dsRBD2 as the CSPs of dsRBD2-0 are higher and show an orientational bias to α1 and the β-sheet (Figure 6a,b). When comparing CSPs of dsRBD2-0 with residues we kept fixed in our designs and proposed to be involved in the DHX9_core_ interaction as predicted by AlphaFold2 and AlphaFold3, we identified both fixed residues that participate in the interaction (A15, G38, I51, Y52, R59) and several that do not (N42, S79, R82). In addition, we identified newly designed sequence regions interspersed with fixed residues which observe a change in the chemical shift upon addition of DHX9_core_, suggesting the formation of larger, newly established interaction sites between dsRBD2-0 and DHX9_core_. These regions include N18-H26, F32-G38, M49-V53 as well as R59-E65 and support the conclusion that dsRBD2-0 primarily interacts with DHX9_core_ through its N-terminal helix and the three β-strands. However, most of the interactions between dsRBD2-0 and DHX9_core_ predicted by AlphaFold3 could not be experimentally confirmed, highlighting the limitations of predictive modeling (Figure 2c). Moreover, dsRBD2-0 inhibits DHX9 activity in a concentration-dependent manner, supporting the proposed DHX9 interface as a druggable target and suggesting that a protein-based approach may be effective in inhibiting DHX9 (Figure 6e). Due to the weak interaction of dsRBD2-0 to DHX9_core_, complete inhibition of DHX9 may not be achieved, as DHX9-dsRBD2 can still engage this interface and sustain helicase function.

In contrast to dsRBD2-0, dsRBD2-3 shows minimal changes in the signal intensity, indicating that it largely tumbles freely in solution and does not extensively interact with DHX9_core_. Consistent with this observation, dsRBD2-3 exhibits weak changes in the chemical shift, whereas the CSPs do not show a specific trend and are distributed across the structure, suggesting rather nonspecific or transient contacts than a defined binding interface (Figure 6c). It is therefore possible that dsRBD2-3 does not bind at the intended interface but instead interacts elsewhere on DHX9_core_, which may explain the lack of inhibitory activity observed specifically for dsRBD2-3 (Figure 6f).

dsRBD5-3 also displays a weak overall interaction with DHX9_core_, as its signal intensity remains identical to dsRBD2-3 (0.92). While the β-strands show little to no CSPs, the α_1_ helix exhibits the strongest CSPs in dsRBD5-3 compared to the remaining sequence (Figure 6d). Notably, despite engaging fewer interaction sites, dsRBD5-3 still inhibits DHX9 helicase activity and follows a concentration-dependent trend similar to dsRBD2-0, suggesting that the interaction through the N-terminal α-helix may be crucial for functional inhibition (Figure 6g).

Unexpectedly, we identified two N- and C-terminal linker residues, H7 and V90, that exhibit high CSPs across all designs. These regions were initially considered irrelevant for dsRBD-core interactions and helicase inhibition. However, our findings suggest that the flanking regions of the engineered dsRBDs may also contribute to interactions with DHX9_core_ and should therefore be considered in future designs. Moreover, the intrinsic flexibility of these residues, arising from their location within disordered regions, may permit adaptable positioning of H7 and V90, enabling optimal orientation towards DHX9_core_ in all designs.

The C-terminal α-helix is not involved in the interaction in any of the designs, confirming MLE and DHX9 structure and prediction. It could therefore be omitted in future designs to reduce size. Future designs do not necessarily need to adopt a dsRBD-fold; a minimal scaffold consisting of α_1_ helix and possibly the three β-strands may be sufficient, provided these elements can form the required contacts and effectively block the interface, thereby preventing dsRBD2 binding and inhibiting helicase activity. Taken together two out of our four experimentally tested dsRBD designs show an interaction with DHX9_core_ and have an inhibitory effect on DHX9 helicase activity comparable to DHX9 dsRBD2^RBM^ (Figure 6h).

## Discussion

DHX9 has emerged as a therapeutic target due to its role in cancer development and tumor progression, with elevated expression levels observed in several cancer types^22,26-33,37-40^. Inhibiting DHX9 has been shown to suppress tumor growth and enhance chemotherapy sensitivity, highlighting its potential as a therapeutic target and for novel cancer therapies^23,33^. Here, we present a novel target site for inhibitors of DHX9 which has not been described before, namely the dsRBD2-core interface. We could show that the interaction between dsRBD2 and DHX9_core_ is crucial for RNA helicase activity. Targeting the specific region of RecA2, OB-fold and HA2 domain is likely specific for DHX9 and distinct from other human helicases as the interaction with the auxiliary domain dsRBD2 seems to be exclusive for DHX9. While small-molecule inhibitors have been developed for DHX9, our protein-based design approach provides a complementary strategy for DHX9 regulation and inhibition and a proof-of-principle from which potent peptide-like inhibitors can be derived. Due to adaptability of cancer cells and patient-dependent variability in therapeutic processes, the development of alternative inhibitory strategies remains essential. Peptide- and protein-derived compounds offer distinct advantages, including high specificity for selected target sites, such as the DHX9_core_, and the ability to engage larger protein surface areas.

Furthermore, peptide- and protein-based drugs harbor low toxicity and immunogenicity^59^. However, cell permeability for larger molecules, such as peptide- and protein-derived compounds, is relatively low due to large size, charge and hydrophilicity^60^. Furthermore, peptide- and protein-based therapeutics suffer from limited bioavailability due to degradation and rapid metabolism^59,61^. This can be overcome by protecting peptides and proteins with polymeric or silk-fibroin based materials^62–64^. Another possibility is the delivery via mRNA. Packaging of the mRNA into lipid nanoparticles protects the mRNA from nuclease degradation, increases its stability in physiological fluids and allows the uptake into cells via micropinocytosis and clathrin-or caveolae-mediated endocytosis^65–67^.This method of packaging offers additional advantages as the nanoparticles can be selectively delivered to specific cell types such as cancer cells by coating of nanoparticles with antibodies or by adjusting the proportions of its lipid components^68–70^.

Our design approach provides practical benefits, including reduced computational demands as we do not generate extensive small molecule libraries but only a small pool of sequences and models. Rather than designing *de novo* sequences, we used dsRBD2 as a template, retaining potentially important residues to enable the generation of structure- and function-based designs that specifically target the DHX9 interface. Furthermore, by redesigning dsRBD2 we may reduce side effects associated with currently unknown cellular functions of dsRBD2 when introducing them as DHX9 inhibitors into cells. The effectiveness of our strategy is supported by the biochemical characterization of our designs as the first round of design generation produced soluble, thermostable, folded DHX9-binding proteins which reduce helicase activity.

Although our designs currently exhibit low affinity, the results demonstrate the potential of both the design strategy and targeting of the dsRBD2-core interface of DHX9. Out of four designs, three were properly folded, with dsRBD2-0 and dsRBD5-3 providing particularly promising starting points for sequence optimization. In a next step, these designs will need to be optimized to achieve higher affinity binding to effectively inhibit DHX9 activity. Notably, dsRBD2-0 and dsRBD5-3 displayed the highest degree of novelty and performed best in our experiments, further highlighting the potential of our design pipeline. Additionally, the size of the designs could be reduced – for example by omitting the α2 helix, which did not appear to be critical for the DHX9_core_ interaction in our designs. Future designs targeting the dsRBD2-core interface may therefore not require the dsRBD-fold, provided that the key interaction features are preserved. Our approach can be extended to other RNA helicases and cancer-associated enzymes. By identifying specific key interaction sites and their specific binder, new binders can be generated targeting the interaction site, thereby inhibiting the protein of interest. Recently developed protein design tools such as RFdiffiusion, BindCraft or AlphaDesign can be used to generate *de novo* protein binders which can be specifically designed to function as novel therapeutics for targeted protein inhibition reaching nanomolar affinities^71–73^. Using BindCraft a broad spectrum of sequences and structures can be generated targeting a desired protein-interface and achieving nanomolar affinities for therapeutic targets such as MDM2 or WDR5^72,74^. In addition, AlphaDesign has also been successfully used to design inhibitors of retron cold anaerobix toxin (RcaT), a bacterial retron-encoded toxin, which is used to defend against viral infection, demonstrating its potential use for designing novel therapeutics^73^. Future protein designs targeting the dsRBD2-core interface in DHX9 may also be generated with alternative protein design tools to create novel folds with optimized binding interfaces and achieve higher binding affinities.

## Material and Methods

### Expression and purification of DHX9 constructs

DHX9 dsRBD2 (aa 167-267) and dsRBD2^RBM^ (aa 167-267) containing the point mutations K182A, N234, K235A and K236A were cloned into pETM11 featuring an N-terminal His-tag and a TEV cleavage site. The plasmids were first transformed into *Escherichia coli* BL21 (DE3). An LB culture was induced at OD_600_ = 0.8 by addition of 1 mM IPTG and subsequently grown overnight at 18 °C. Cells were harvested for 15 min at 8983 x g and 4 °C and pellets were resuspended in lysis buffer (50 mM NaPi, 500 mM NaCl, 20 mM imidazole, 2 mM MgCl_2_, 1 mM DTT, pH 7.5) supplemented with benzonase and protease inhibitor cocktail (Roche). Cell lysis was done using a microfluidizer and the lysate was pelleted at 27216 x g and 4 °C for 30 min. The supernatant was loaded onto a HisTrap HP column (Cytiva), washed with wash buffer (50 mM NaPi, 500 mM NaCl, 20 mM Imidazole, 1 mM DTT, pH 7.5) and eluted in a linear imidazole gradient in buffer containing 50 mM NaPi, 500 mM NaCl, 500 mM imidazole, 1 mM DTT, pH 7.5. The His-tag was removed during dialysis into 20 mM NaPi, 50 mM NaCl, 1 mM DTT, pH 6.5 overnight at 4 °C by addition of 30 nmol TEV protease. The tag was removed by loading the sample onto the HisTrap HP column and protein containing fractions were further applied to a 1 mL HiTrap Heparin HP column to remove bound nucleic acids in the case of DHX9 dsRBD2. Finally, protein containing fractions were loaded onto a size-exclusion chromatography on the HiLoad 26/600 Superdex S75 pg column equilibrated in 20 mM NaPi, 50 mM NaCl, 1 mM DTT, pH 6.5.

DHX9_core_ was cloned into pFastBac-HTA containing a N-terminal His-tag with TEV cleavage site and expressed in High5 insect cells using recombinant baculovirus produced in Sf9 cells. After infection of 1.2 x 10^6^ cells/mL High5 cells in a 1:100 ratio with the baculovirus, DHX9_core_ was expressed for 72 h. Cells were harvested at 900 x g and 4 °C for 30 min and the pellet was resuspended in lysis buffer (20 mM HEPES, 500 mM NaCl, 5% glycerol, 20 mM imidazole, 2 mM MgCl_2_, 0.005% Triton X-100, 1 mM DTT, pH 7.5) supplemented with benzonase and protease inhibitor cocktail (Roche). Cells were lysed in a douncer and pelleted at 27216 x g and 4 °C for 30 min. The supernatant was rotated for 2 h at 4 °C with 1 mL Ni-NTA agarose (Qiagen) equilibrated in wash buffer (20 mM HEPES, 500 mM NaCl, 5% glycerol, 20 mM Imidazole, 1 mM DTT, pH 7.5). Beads were pelleted at 233 x g for 2 min and the supernatant was removed. Beads were washed with wash buffer and protein was eluted with 20 mM HEPES, 500 mM NaCl, 5% glycerol, 500 mM imidazole, 1 mM DTT, pH 7.5. The His-tag was removed during dialysis into 20 mM HEPES, 200 mM NaCl, 1 mM DTT, pH 7.5 by addition of 30 nmol TEV protease. Cleaved protein was further purified by a reverse IMAC followed by cation exchange chromatography. The sample was loaded onto a 1 mL HiTrap Heparin HP column, washed with 20 CV low salt buffer (20 mM HEPES, 200 mM NaCl, 1 mM DTT, pH 7.5) and eluted in 20 CV with a linear gradient of high salt buffer (20 mM HEPES, 1 M NaCl, 1 mM DTT, pH 7.5). Pooled fractions were loaded onto a HiLoad 16/600 Superdex S200 pg column equilibrated in 20 mM HEPES, 200 mM NaCl, 1 mM DTT, pH 7.5.

All proteins were frozen in liquid nitrogen and stored at −80 °C.

### Computational approach of redesigning DHX9 dsRBD2

The sequences of the designed dsRBDs were generated using ProteinMPNN^47^. Briefly, the sequence alignment of DHX9 and MLE’s dsRBDs presented in Jagtap et. al (2019) was used as reference for selection of fixed residues^75^. In six design rounds, 30 sequences were generated with ProteinMPNN with following settings: *omitted aa: M+C, sampling temperature = 0*.*1, backbone noise = 0*.*1, model: vanilla-v_48_020*. Subsequently the sequences were predicted using ColabFold^48,49^ (default settings). Models with a global pLDDT score ≥ 80 were selected for further analysis. To analyze the DHX9-dsRBD complex, the selected sequences were predicted with AlphaFold3^50^. The complexes were further evaluated based on the pAE and local pLDDT. In parallel, the selected sequences were docked using the HADDOCK Web Server^51,52^. Final complexes were analyzed according to buried surface area (BSA), cluster RMSD and HADDOCK scores. The models of the complexes in both methods were further structurally evaluated by their interface and the predicted protein-protein interactions. These models were superimposed with the structure of MLE in the dsRBD2 bound state (PDB 8PJB) to analyze the domain orientation of the designed dsRBDs^46^. In the final filtering step, the selected sequences were predicted regarding the potential toxicity and solubility for recombinant expression in *Escherichia coli* using the Web Server CSM-Toxin^54^ and ProteinSol^55^. In the end, the chosen sequences were tested regarding their novelty using the Web Server SeqNovelty (*Blast nr. of sequences: 500, align mode: pairwise, identity cutoff: 0*.*3, open gap penalty: −10, extend gap penalty: −0*.*5*)^56^.

### Cloning, expression and purification of dsRBD designs

Gene fragments were ordered from Azenta and cloned into the N-terminal Thioredoxin A, His_6_-Tag and TEV protease containing expression vector pETM20. The designs were expressed in *Escherichia coli* BL21 (DE3) by addition of 1 mM IPTG at OD_600_ = 0.8 and overnight growth at 18 °C. The cells were harvested and resuspended in lysis buffer (50 mM NaPi, 500 mM NaCl, 1 M urea, 1 mM DTT, pH 7.5) supplemented with benzonase and protease inhibitor cocktail (Roche). Next, cell lysis was done using a fluidizer and the lysate was pelleted at 27216 x g and 4 °C for 30 min. The supernatant was loaded onto a HisTrap HP column (Cytiva), washed with wash buffer (50 mM NaPi, 500 mM NaCl, 1 mM DTT, pH 7.5) and eluted with a linear imidazole gradient in buffer containing 50 mM NaPi, 500 mM NaCl, 500 mM imidazole, 1 mM

DTT, pH 7.5. The Thioredoxin A-His_6_-tag was removed during dialysis into 20 mM NaPi, 100 mM NaCl, 1 mM DTT, pH 7.0 overnight by addition of 30 nmol TEV protease. Next, the tag was removed by loading the sample onto the HisTrap HP column and the flow through was further applied onto a size-exclusion chromatography on the HiLoad 26/600 Superdex S75 pg column equilibrated in NMR buffer (20 mM NaPi, 100 mM NaCl, 1 mM DTT, pH 7.0).

For NMR experiments, the proteins were expressed in M9 minimal media containing 1.5 g/l ^13^C-labelled glucose and/or 0.5 g/l ^15^N-labelled ammonium sulfate. NMR samples were prepared in 10% D_2_O for the deuterium lock.

### Dynamic light scattering and melting temperature determination

Hydrodynamic diameters and melting temperatures were determined using Uncle (UNchained Labs, Pleasanton, CA). All samples were measured in triplicates at 1.5 mg/ml concentrations with 9 µl loaded into each capillary.

The hydrodynamic diameter was determined by dynamic light scattering. After an incubation time of 180 s the samples were measured at 660 nm with 4 acquisitions and an acquisition time of 5 s. To determine the melting temperatures, differential scanning fluorimetry was performed with the same samples in a temperature ramp ranging from 20 °C to 95 °C at 0.6 °C/min after an incubation time of 180 s. Fluorescence spectra were collected from 300 nm to 430 nm. Melting temperatures were determined from the inflection point of the barycentric mean.

### Circular dichroism

CD spectra were collected with a Jasco J-1100 CD spectrometer. All experiments were recorded in 20 mM NaP_i_, 50 mM NaCl and 1 mM DTT using a protein concentration of 0.2 mg/ml. Far-ultraviolet-CD spectra were recorded from 190-260 nm at 25 °C in a 1 mm cuvette with a 1 nm bandwidth and scanning rate of 50 nm/min. For each protein 20 spectra were accumulated and averaged. Data were buffer corrected and normalized to mean residue molar ellipticity using following equation: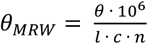, where θis in mdeg, path length l is in mm, concentration c is in µM, and the number of peptide bonds is n.

### Fluorescence polarization assays

Fluorescence polarization assays were carried out in 20 mM NaPi, 50 mM NaCl, 0.005% Igepal CA-630,1 mM DTT, pH 6.5 and with 6-Carboxyfluorescein (6-FAM) labeled RNA. ssRNA (5’-UUUUUUUUUUUU-6-FAM 3’) was labelled at the 3’ end with 6FAM. dsRNA was annealed by heating up SL7mod_up (5’ GUGUAAAAUGUUGCUAGCA 3’) and SL7mod_down (5’ 6-FAM-UGCUAGUAACGUUUUACGCCC 3’) at 95 °C for 30 s and subsequent incubation on ice for 20 min. All RNAs were purchased from Biomers. A serial dilution series of protein was prepared in 2x buffer in a volume of 15 µ in Corning 384-well plates. Addition of 15 µl of a 5 nM RNA stock solution to each well resulted in a total reaction volume of 30 µl and a final RNA concentration of 2.5 nM. The reaction was incubated for 30 min in the dark at room temperature. Fluorescence polarization was measured with the Synergy 2 microplate reader from BioTek with excitation and emission wavelengths of 485 nm and 528 nm, respectively. Data were plotted in Graph Prism v9 and were fit to a Sigmoidal 4PL equation, in which X is log(concentration). To calculate IC50 values 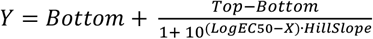 was applied, where Top and Bottom are upper and lower boundaries of signals.

### Helicase assay

Helicase assays were carried out in 20 mM Tris, 100 mM NaCl, 5 mM MgCl2, 0.005% IGEPAL CA-630, 2 mM DTT, pH 7.5 with dsRNA containing a BHQ-1 and 6-FAM quencher-dye pair. SL7-up_BHQ1 (5’ GUGUAAAAUGUUGCUAGCA-BHQ1 3’) and 6FAM_SL7-down (5’ 6-FAM-UGCUAGUAACGUUUUACGCCCUCUUUCUUUCUU 3’) were mixed in equimolar ratio at 1 µM in 20 mM Tris, 100 mM NaCl, pH 7.4, heated to 95 ºC for 1 min and then slowly cooled down to 4 ºC in 5 °C/30 s steps using a PCR cycler. A SL7-up DNA competitor was added as a competitor to prevent re-annealing of the displaced SL7-up_BHQ1 strand to 6FAM_SL7-down upon helicase unwinding activity at 25-fold molar excess. As negative controls, reactions without protein and as positive controls, reactions lacking SL7-up_BHQ1 were measured. The assays were carried out in 30 µl reaction volumes with 300 nM protein and a final concentration of 5 nM annealed RNA in 384-well plates. A dilution series of the designs, which were dialyzed into 20 mM Tris, 100 mM NaCl, 1 mM DTT, pH 7.5, was prepared from 30 µM to 1.875 µM in the wells. Samples were prepared identically to maintain comparability. Reactions were incubated for 20 min at room temperature in the dark. Baseline was measured for 3 min and the time point of ATP addition was set to zero. All helicase assays were performed in duplicates.

### NMR data acquisition and structure determination

All spectra were recorded using Avance IIIHD Bruker NMR spectrometers with proton Larmor resonance frequencies of 700 MHz, 900 MHz and 1000 MHz at 298 K, equipped with cryogenically cooled triple resonance gradient probes.

For protein resonance assignment standard double and triple resonance through-bond experiments were recorded^76. 13^C- and ^15^N edited NOESY spectra (mixing time 120 ms) were performed to obtain distance restraints. Apodization weighted sampling was used for acquisition of triple resonance spectra^77^, and non-uniform sampling was applied for ^15^N-edited NOESY spectra. NMR data were processed using NMPipe^78^ or in-house routines and analyzed with Cara (https://cara.nmr.ch) or NMRViewJ^79^.

Distance restraints for structure calculation were derived from ^15^N-edited NOESY and ^13^C-edited NOESY spectra. NOESY cross peaks were classified according to their relative intensities and converted to distance restraints with upper limits of 3.0 Å (strong), 4.0 Å (medium), 5.0 Å (weak), and 6.0 Å (very weak). For ambiguous distance restraints the r^-6^ summation over all assigned possibilities defined the upper limit. Dihedral angle restraints were derived from backbone chemical shifts using TALOS+^80^. Hydrogen bonds were included for backbone amide protons in regular secondary structure, when the amide proton does not show a water exchange cross peak in the ^15^N-edited NOESY spectrum.

The structure calculations were performed with the program XPLOR-NIH 1.2.1^81^ using a three-step simulated annealing protocol with floating assignment of prochiral groups including a conformational database potential. The 20 (out of total 80) structures showing the lowest values of the target function excluding the database potential were further analyzed with X-PLOR^81^, PyMOL, and PROCHECK 3.5.4^82^.

### NMR titration experiments

NMR titration experiments were performed at 10 µM protein concentration in 20 mM NaP_i_, 50 mM NaCl, 1 mM DTT, pH 6.5 and 10% D_2_O. ^15^N labelled proteins were titrated to a 1:1 ratio of DHX9_core_, which was dialyzed into the same buffer, and monitored by recording ^1^H, ^15^N HSQC spectra. Spectra were processed using NMRPipe^78^. Data analysis for calculating chemical shift perturbations (CSPs) and signal-intensity-ratios were performed using Sparky^83^. CSPs were calculated using the chemical shifts of the free state and the bound state using the following equation: 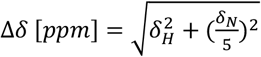. The significance threshold of the CSPs was determined as previously described^84^. Secondary chemical shifts were calculated by subtracting the chemical shifts δ(Cα) and δ(Cβ) of a random coil from δ(Cα) or δ(Cβ) of the respective amino acid resulting in Δδ(Cα) and Δδ(Cβ). Followed by subtracting Δδ(Cβ) from Δδ(Cα). Positive Δδ(Cα)-Δδ(Cβ) values indicate α-helical propensity, whereas negative values indicate β-sheet propensity^58,57^.

## Data availability

Atomic coordinates of the NMR solution structure of dsRBD2-0 was deposited to the Protein Data Bank (PDB) under accession number 28PM. Chemical shifts and NMR data for structure determination of dsRBD2-0 were deposited to the BMRB under accession number 35035. Backbone assignments of DHX9-dsRBD2, dsRBD2-3 and dsRBD5-3 were deposited to the BMRB with accession codes 53569, 53568 and 53567, respectively.

## Acknowledgements

J.H. gratefully acknowledges support by the European Molecular Biology Laboratory.

## Author contributions

Conceptualization, N.L., E.F., and J.H.; methodology: N.L. (NMR, biochemical and biophysical characterization, protein design), E.F. (protein design), K.S. (NMR); formal analysi:, N.L., L.H. (NMR), K.S. (NMR) and J.H. (NMR); investigation: N.L. and E.F.; writing – original draft: N.L.; writing – review & editing: N.L., E.F., K.S. and J.H.; project administration: N.L., J.H.; funding acquisition: J.H.; resources: J.H.; supervision: J.H.

## Additional information

Supplementary information

## References

1. Dakal, T. C. et al. Oncogenes and tumor suppressor genes: functions and roles in cancers. MedComm 5, e582 (2024).

2. Zhang, S. et al. Tumor initiation and early tumorigenesis: molecular mechanisms and interventional targets. Signal Transduct. Target. Ther. 9, 149 (2024).

3. Wang, W.-J., Li, L.-Y. & Cui, J.-W. Chromosome structural variation in tumorigenesis: mechanisms of formation and carcinogenesis. Epigenetics Chromatin 13, 49 (2020).

4. Obeng, E. A., Stewart, C. & Abdel-Wahab, O. Altered RNA Processing in Cancer Pathogenesis and Therapy. Cancer Discov. 9, 1493–1510 (2019).

5. Jia, X. et al. Protein translation: biological processes and therapeutic strategies for human diseases. Signal Transduct. Target. Ther. 9, 44 (2024).

6. Heerma van Voss, M. R., van Diest, P. J. & Raman, V. Targeting RNA helicases in cancer: The translation trap. Biochim. Biophys. Acta BBA - Rev. Cancer 1868, 510–520 (2017).

7. Cai, W. et al. Wanted DEAD/H or Alive: Helicases Winding Up in Cancers. JNCI J. Natl. Cancer Inst. 109, djw278 (2017).

8. Bourgeois, C. F., Mortreux, F. & Auboeuf, D. The multiple functions of RNA helicases as drivers and regulators of gene expression. Nat. Rev. Mol. Cell Biol. 17, 426–438 (2016).

9. Lang, N., Jagtap, P. K. A. & Hennig, J. Regulation and mechanisms of action of RNA helicases. RNA Biol. 21, 24–38 (2024).

10. Bohnsack, K. E., Yi, S., Venus, S., Jankowsky, E. & Bohnsack, M. T. Cellular functions of eukaryotic RNA helicases and their links to human diseases. Nat. Rev. Mol. Cell Biol. 1–21 (2023) doi:10.1038/s41580-023-00628-5.

11. Arna, A. B. et al. Synthetic lethal interactions of DEAD/H-box helicases as targets for cancer therapy. Front. Oncol. 12, 1087989 (2022).

12. Andrisani, O. et al. Biological functions of DEAD/DEAH-box RNA helicases in health and disease. Nat. Immunol. 23, 354–357 (2022).

13. Calame, D. G. et al. Monoallelic variation in DHX9, the gene encoding the DExH-box helicase DHX9, underlies neurodevelopment disorders and Charcot-Marie-Tooth disease. Am. J. Hum. Genet. 110, 1394–1413 (2023).

14. Jankowsky, E. RNA helicases at work: binding and rearranging. Trends Biochem. Sci. 36, 19–29 (2011).

15. Bleichert, F. & Baserga, S. J. The Long Unwinding Road of RNA Helicases. Mol. Cell 27, 339–352 (2007).

16. Fairman-Williams, M. E., Guenther, U.-P. & Jankowsky, E. SF1 and SF2 helicases: family matters. Curr. Opin. Struct. Biol. 20, 313–324 (2010).

17. Lee, C. G. & Hurwitz, J. A new RNA helicase isolated from HeLa cells that catalytically translocates in the 3’ to 5’ direction. J. Biol. Chem. 267, 4398–4407 (1992).

18. Zhang, S. & Grosse, F. Multiple functions of nuclear DNA helicase II (RNA helicase A) in nucleic acid metabolism. Acta Biochim. Biophys. Sin. 36, 177–183 (2004).

19. Zhang, S. & Grosse, F. Nuclear DNA helicase II unwinds both DNA and RNA. Biochemistry 33, 3906–3912 (1994).

20. Chakraborty, P. & Grosse, F. Human DHX9 helicase preferentially unwinds RNA-containing displacement loops (R-loops) and G-quadruplexes. DNA Repair 10, 654–665 (2011).

21. Jain, A. et al. DHX9 helicase is involved in preventing genomic instability induced by alternatively structured DNA in human cells. Nucleic Acids Res. 41, 10345–10357 (2013).

22. Ding, X. et al. A DHX9-lncRNA-MDM2 interaction regulates cell invasion and angiogenesis of cervical cancer. Cell Death Differ. 26, 1750–1765 (2019).

23. Lee, T. & Pelletier, J. The biology of DHX9 and its potential as a therapeutic target. Oncotarget 7, 42716–42739 (2016).

24. Aktaş, T. et al. DHX9 suppresses RNA processing defects originating from the Alu invasion of the human genome. Nature 544, 115–119 (2017).

25. Zhou, Y. et al. RNA damage compartmentalization by DHX9 stress granules. Cell 187, 1701–1718.e28 (2024).

26. Gulliver, C., Hoffmann, R. & Baillie, G. S. The Enigmatic Helicase DHX9 and Its Association With the Hallmarks of Cancer. Future Sci. OA 7, FSO650 (2021).

27. Schlegel, B. P., Starita, L. M. & Parvin, J. D. Overexpression of a protein fragment of RNA helicase A causes inhibition of endogenous BRCA1 function and defects in ploidy and cytokinesis in mammary epithelial cells. Oncogene 22, 983–991 (2003).

28. Guénard, F., Labrie, Y., Ouellette, G., Joly Beauparlant, C. & Durocher, F. Genetic sequence variations of BRCA1-interacting genes AURKA, BAP1, BARD1 and DHX9 in French Canadian Families with high risk of breast cancer. J. Hum. Genet. 54, 152–161 (2009).

29. Huang, N. et al. DHX9-mediated pathway contributes to the malignant phenotype of myelodysplastic syndromes. iScience 26, 106962 (2023).

30. Chen, X. et al. High Levels of DEAH-Box Helicases Relate to Poor Prognosis and Reduction of DHX9 Improves Radiosensitivity of Hepatocellular Carcinoma. Front. Oncol. 12, (2022).

31. Chellini, L., Pieraccioli, M., Sette, C. & Paronetto, M. P. The DNA/RNA helicase DHX9 contributes to the transcriptional program of the androgen receptor in prostate cancer. J. Exp. Clin. Cancer Res. 41, 178 (2022).

32. Liu, S. et al. DHX9 contributes to the malignant phenotypes of colorectal cancer via activating NF-κB signaling pathway. Cell. Mol. Life Sci. 78, 8261–8281 (2021).

33. Cao, S. et al. RNA helicase DHX9 may be a therapeutic target in lung cancer and inhibited by enoxacin. Am. J. Transl. Res. 9, 674–682 (2017).

34. Wang, Y. et al. A pan-cancer analysis of the expression and molecular mechanism of DHX9 in human cancers. Front. Pharmacol. 14, (2023).

35. Mi, J. et al. In Vivo Selection Against Human Colorectal Cancer Xenografts Identifies an Aptamer That Targets RNA Helicase Protein DHX9. Mol. Ther. - Nucleic Acids 5, e315 (2016).

36. Fidaleo, M. et al. Genotoxic stress inhibits Ewing sarcoma cell growth by modulating alternative pre-mRNA processing of the RNA helicase DHX9. Oncotarget 6, 31740–31757 (2015).

37. Frezza, V. et al. DHX9 helicase impacts on splicing decisions by modulating U2 snRNP recruitment in Ewing sarcoma cells. Nucleic Acids Res. 53, gkaf068 (2025).

38. Hong, H. et al. Bidirectional regulation of adenosine-to-inosine (A-to-I) RNA editing by DEAH box helicase 9 (DHX9) in cancer. Nucleic Acids Res. 46, 7953–7969 (2018).

39. Liu, H. et al. Single-cell transcriptomics reveal DHX9 in mature B cell as a dynamic network biomarker before lymph node metastasis in CRC. Mol. Ther. - Oncolytics 22, 495–506 (2021).

40. Chellini, L., Pieraccioli, M., Sette, C. & Paronetto, M. P. The DNA/RNA helicase DHX9 contributes to the transcriptional program of the androgen receptor in prostate cancer. J. Exp. Clin. Cancer Res. 41, 178 (2022).

41. Nogales, V. et al. Epigenetic inactivation of the putative DNA/RNA helicase SLFN11 in human cancer confers resistance to platinum drugs. Oncotarget 7, 3084–3097 (2015).

42. Frezza, V. et al. DHX9 helicase impacts on splicing decisions by modulating U2 snRNP recruitment in Ewing sarcoma cells. Nucleic Acids Res. 53, gkaf068 (2025).

43. Lee, T., Paquet, M., Larsson, O. & Pelletier, J. Tumor Cell Survival Dependence on the DHX9 DExH-Box Helicase. Oncogene 35, 5093–5105 (2016).

44. Daniels, M. H. et al. Discovery of ATX968: An Orally Available Allosteric Inhibitor of DHX9. J. Med. Chem. https://doi.org/10.1021/acs.jmedchem.5c00252 (2025) doi:10.1021/acs.jmedchem.5c00252.

45. Prabu, J. R. et al. Structure of the RNA Helicase MLE Reveals the Molecular Mechanisms for Uridine Specificity and RNA-ATP Coupling. Mol. Cell 60, 487–499 (2015).

46. Jagtap, P. K. A. et al. Structural basis of RNA-induced autoregulation of the DExH-type RNA helicase maleless. Mol. Cell 83, 4318–4333.e10 (2023).

47. Dauparas, J. et al. Robust deep learning–based protein sequence design using ProteinMPNN. Science 378, 49–56 (2022).

48. Jumper, J. et al. Highly accurate protein structure prediction with AlphaFold. Nature 596, 583–589 (2021).

49. Google Colab. https://colab.research.google.com/github/sokrypton/ColabFold/blob/main/AlphaFold2.ipynb.

50. Abramson, J. et al. Accurate structure prediction of biomolecular interactions with AlphaFold 3. Nature 630, 493–500 (2024).

51. Dominguez, C., Boelens, R. & Bonvin, A. M. J. J. HADDOCK: A Protein-Protein Docking Approach Based on Biochemical or Biophysical Information. J. Am. Chem. Soc. 125, 1731–1737 (2003).

52. HADDOCK2.4 small molecule binding site screening. Bonvin Lab https://www.bonvinlab.org/education/HADDOCK24/HADDOCK24-binding-sites/.

53. Fu, Q. & Yuan, Y. A. Structural insights into RISC assembly facilitated by dsRNA-binding domains of human RNA helicase A (DHX9). Nucleic Acids Res. 41, 3457–3470 (2013).

54. Morozov, V., Rodrigues, C. H. M. & Ascher, D. B. CSM-Toxin: A Web-Server for Predicting Protein Toxicity. Pharmaceutics 15, 431 (2023).

55. Hebditch, M., Carballo-Amador, M. A., Charonis, S., Curtis, R. & Warwicker, J. Protein– Sol: a web tool for predicting protein solubility from sequence. Bioinformatics 33, 3098–3100 (2017).

56. SeqNovelty. https://fuerstlab.shinyapps.io/SeqNovelty/.

57. Wishart, D. S. & Sykes, B. D. The 13C Chemical-Shift Index: A simple method for the identification of protein secondary structure using 13C chemical-shift data. J. Biomol. NMR 4, 171–180 (1994).

58. Wishart, D. S., Bigam, C. G., Holm, A., Hodges, R. S. & Sykes, B. D. 1H, 13C and 15N random coil NMR chemical shifts of the common amino acids. I. Investigations of nearest-neighbor effects. J. Biomol. NMR 5, 67–81 (1995).

59. Bruno, B. J., Miller, G. D. & Lim, C. S. Basics and recent advances in peptide and protein drug delivery. Ther. Deliv. 4, 1443–1467 (2013).

60. Muheem, A. et al. A review on the strategies for oral delivery of proteins and peptides and their clinical perspectives. Saudi Pharm. J. SPJ Off. Publ. Saudi Pharm. Soc. 24, 413–428 (2016).

61. Shaji, J. & Patole, V. Protein and Peptide drug delivery: oral approaches. Indian J. Pharm. Sci. 70, 269–277 (2008).

62. Liechty, W. B., Kryscio, D. R., Slaughter, B. V. & Peppas, N. A. Polymers for drug delivery systems. Annu. Rev. Chem. Biomol. Eng. 1, 149–173 (2010).

63. Zhao, H. et al. Polymer-based nanoparticles for protein delivery: design, strategies and applications. J. Mater. Chem. B 4, 4060–4071 (2016).

64. Wu, J., Sahoo, J. K., Li, Y., Xu, Q. & Kaplan, D. L. Challenges in delivering therapeutic peptides and proteins: A silk-based solution. J. Controlled Release 345, 176–189 (2022).

65. Kim, J., Eygeris, Y., Gupta, M. & Sahay, G. Self-assembled mRNA vaccines. Adv. Drug Deliv. Rev. 170, 83–112 (2021).

66. Blakney, A. K., McKay, P. F., Yus, B. I., Aldon, Y. & Shattock, R. J. Inside out: optimization of lipid nanoparticle formulations for exterior complexation and in vivo delivery of saRNA. Gene Ther. 26, 363–372 (2019).

67. Li, B., Zhang, X. & Dong, Y. Nanoscale platforms for messenger RNA delivery. Wiley Interdiscip. Rev. Nanomed. Nanobiotechnol. 11, e1530 (2019).

68. Kranz, L. M. et al. Systemic RNA delivery to dendritic cells exploits antiviral defence for cancer immunotherapy. Nature 534, 396–401 (2016).

69. Krienke, C. et al. A noninflammatory mRNA vaccine for treatment of experimental autoimmune encephalomyelitis. Science 371, 145–153 (2021).

70. Kedmi, R. et al. A modular platform for targeted RNAi therapeutics. Nat. Nanotechnol. 13, 214–219 (2018).

71. Watson, J. L. et al. De novo design of protein structure and function with RFdiffusion. Nature 620, 1089–1100 (2023).

72. Pacesa, M. et al. One-shot design of functional protein binders with BindCraft. Nature 646, 483–492 (2025).

73. Jendrusch, M. A. et al. AlphaDesign: a de novo protein design framework based on AlphaFold. Mol. Syst. Biol. 21, 1166–1189 (2025).

74. Filius, M. et al. Evaluating BindCraft for Generative Design of High-Affinity Peptides. ACS Chem. Biol. 20, 2991–2998 (2025).

75. Ankush Jagtap, P. K. et al. Structure, dynamics and roX2-lncRNA binding of tandem double-stranded RNA binding domains dsRBD1,2 of Drosophila helicase Maleless. Nucleic Acids Res. 47, 4319–4333 (2019).

76. Sattler, M., Schleucher, J. & Griesinger, C. Heteronuclear multidimensional NMR experiments for the structure determination of proteins in solution employing pulsed field gradients. Prog. Nucl. Magn. Reson. Spectrosc. 34, 93–158 (1999).

77. Simon, B. & Köstler, H. Improving the sensitivity of FT-NMR spectroscopy by apodization weighted sampling. J. Biomol. NMR 73, 155–165 (2019).

78. Delaglio, F. et al. NMRPipe: a multidimensional spectral processing system based on UNIX pipes. J. Biomol. NMR 6, 277–293 (1995).

79. Johnson, B. A. & Blevins, R. A. NMR View: A computer program for the visualization and analysis of NMR data. J. Biomol. NMR 4, 603–614 (1994).

80. Shen, Y., Delaglio, F., Cornilescu, G. & Bax, A. TALOS+: a hybrid method for predicting protein backbone torsion angles from NMR chemical shifts. J. Biomol. NMR 44, 213–223 (2009).

81. Schwieters, C. D., Kuszewski, J. J., Tjandra, N. & Clore, G. M. The Xplor-NIH NMR molecular structure determination package. J. Magn. Reson. 160, 65–73 (2003).

82. Laskowski, R. A., MacArthur, M. W., Moss, D. S. & Thornton, J. M. PROCHECK: a program to check the stereochemical quality of protein structures. J. Appl. Crystallogr. 26, 283–291 (1993).

83. Lee, W., Tonelli, M. & Markley, J. L. NMRFAM-SPARKY: enhanced software for biomolecular NMR spectroscopy. Bioinformatics 31, 1325–1327 (2015).

84. Schumann, F. H. et al. Combined chemical shift changes and amino acid specific chemical shift mapping of protein–protein interactions. J. Biomol. NMR 39, 275–289 (2007).

