## Supplementary Information for "dsRBD Redesign: A Targeted Strategy for Inhibition of RNA Helicase DHX9"

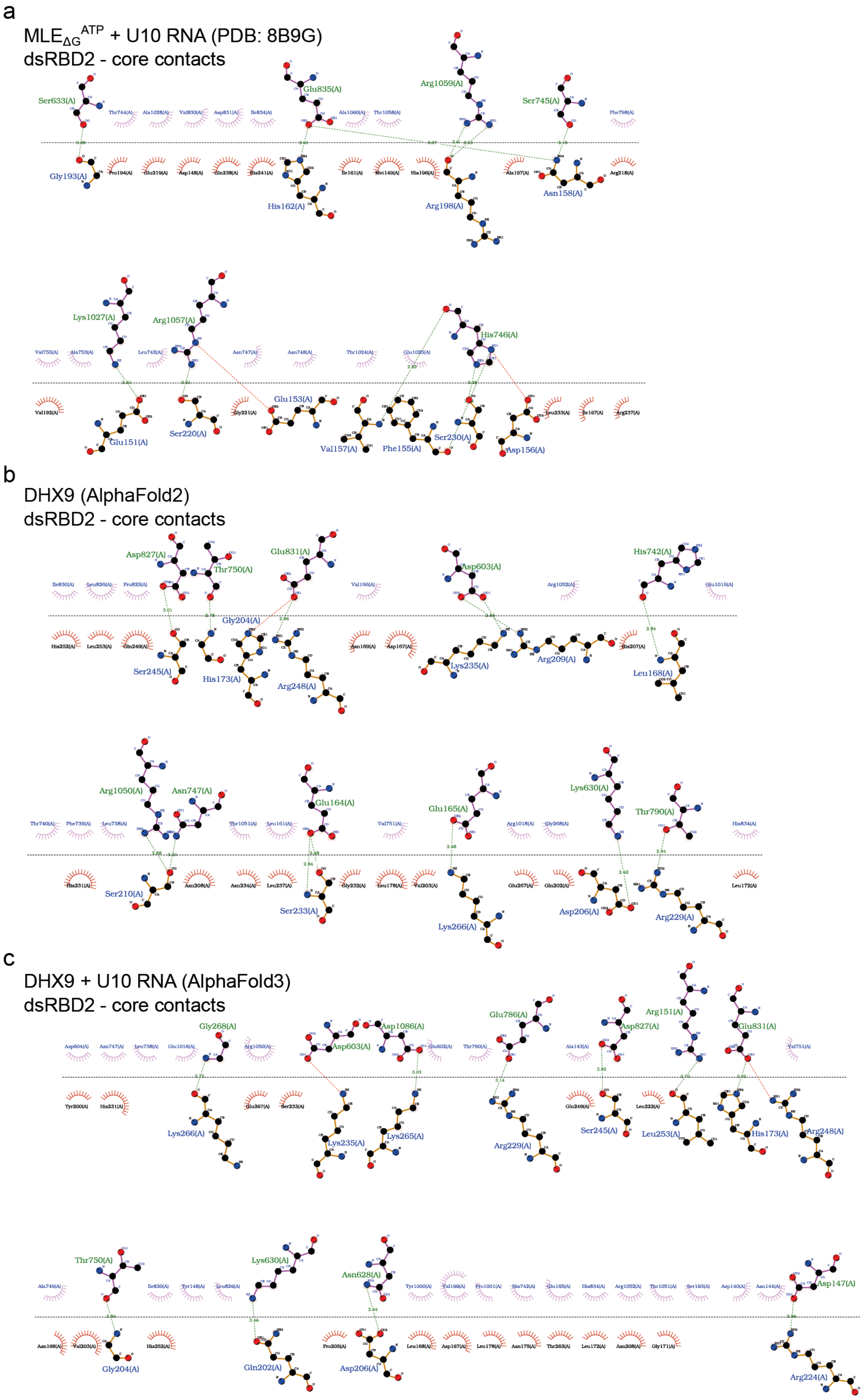


**Figure S1: MLE-dsRBD2 and predicted DHX9-dsRBD2 interactions with the helicase core determined with LigPlot+. (a)** MLE-dsRBD2 contacts with the helicase core in the cryo EM structure of $MLE_{\Delta G}$ in complex with U10 RNA (PDB: 8B9G). **(b)** DHX9-dsRBD2 contacts with the helicase core predicted by the AlphaFold2 model of DHX9.

**(c)** DHX9-dsRBD2 contacts with the helicase core predicted by the AlphaFold3 model of DHX9 in complex with U10 RNA.

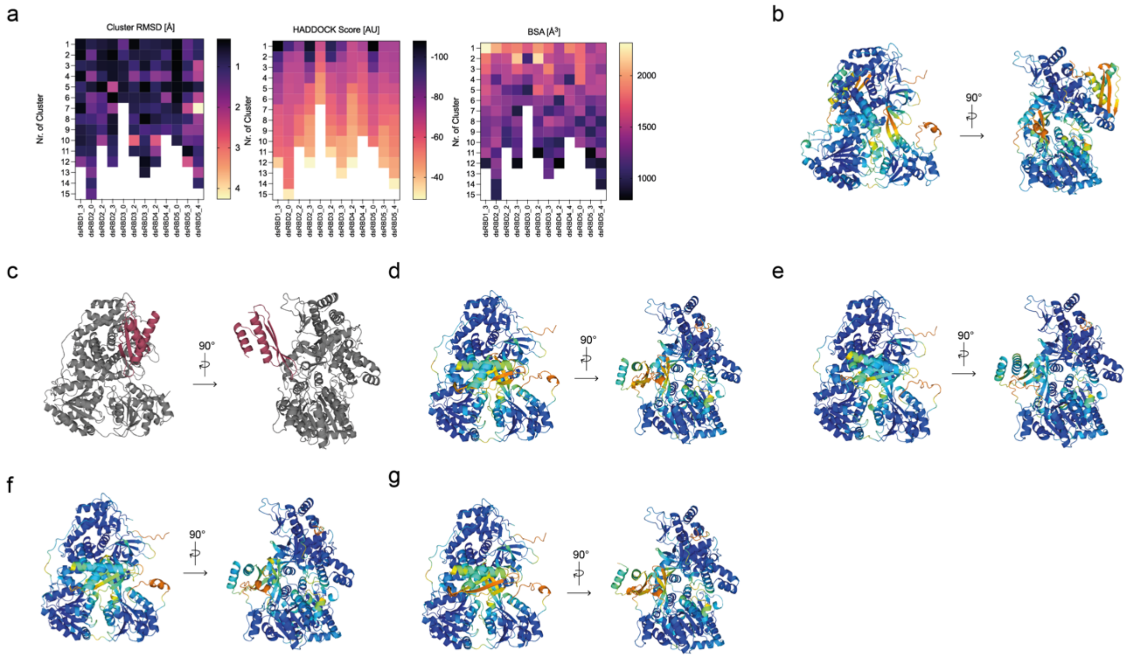

**Figure S2: dsRBD design AlphaFold3 complex predictions and HADDOCK scores. (a)** HADDOCK heat maps of the 12 designs that passed our pLLDT standards (≥ 80) including dsRBD1-3, dsRBD2-0, dsRBD2-3, dsRBD5-3 as well as other not experimentally analyzed designs **(b)** dsRBD4-4, which is not predicted by AlphaFold3 to bind at the dsRBD2-core interface, is shown as a negative example. Colored according to plDDT values **(c)** Negative example of HADDOCK docking illustrated with dsRBD3-2 (red), which does not properly bind to the dsRBD2-core interface. **(d-g)** AlphaFold3 complex predictions of dsRBD1-3 (d), dsRBD2-0 (e), dsRBD2-3 (f), dsRBD5-3 (g).

**Table S1: Computational statistics of the remaining dsRBD designs that passed our pLDDT standards (≥ 80).** [a]AlphaFold3 interface predicted template modeling score. [b]HADDOCK RMSD measuring the distance between the interface backbone atoms and the reference structure. [c]HADDOCK buried surface area. [d]according to lowest HADDOCK score.

| | dsRBD2-2 | dsRBD3-0 | dsRBD3-2 | dsRBD3-3 |
|---|---|---|---|---|
| SeqNovelty | 48 % | 45 % | 48 % | 48 % |
| Fixed residues | 24 % | 24 % | 24 % | 24 % |
| De novo residues | 62 % | 59 % | 62 % | 62 % |
| Conserved residues | 28 % | 30 % | 28 % | 28 % |
| AF3 iptm[a] | 0.57 | 0.50 | 0.52 | 0.52 |
| Haddock-score [AU] | -81 | -67.6 | -98.3 | -87.5 |
| Haddock RMSD [Å][b] | 0.64 | 1.31 | 0.40 | 0.76 |
| Haddock BSA [$Å^3$][c] | 1793.8 | 1655.6 | 1933.2 | 1763.9 |
| No. of clusters | 10 | 6 | 11 | 13 |
| No. of structures in best cluster[d] | 42 | 89 | 43 | 37 |
| CSM Toxin | Non-toxic | Non-toxic | Non-toxic | Non-toxic |
| Protein Sol | 93.7 % | 78.3 % | 89.6 % | 100 % |

| | dsRBD4-2 | dsRBD4-4 | dsRBD5-0 | dsRBD5-4 |
|---|---|---|---|---|
| SeqNovelty | 48 % | 45 % | 48 % | 48 % |
| Fixed residues | 24 % | 24 % | 24 % | 24 % |
| De novo residues | 62 % | 59 % | 62 % | 62 % |
| Conserved residues | 28 % | 30 % | 28 % | 28 % |
| AF3 iptm[a] | 0.32 | 0.15 | 0.42 | 0.57 |
| Haddock-score [AU] | -68.6 | -74.6 | -107.3 | -74.4 |
| Haddock RMSD [Å][b] | 0.69 | 0.63 | 0.43 | 0.89 |
| Haddock BSA [$Å^3$][c] | 1444.2 | 1867.4 | 1752.5 | 1513.6 |
| No. of clusters | 12 | 9 | 10 | 14 |
| No. of structures in best cluster[d] | 43 | 74 | 95 | 24 |
| CSM Toxin | Non-toxic | toxic | Non-toxic | Non-toxic |
| Protein Sol | 88.9 % | 79.7 % | 96.2 % | 87.7 % |
